# Sex-dimorphic gene regulation in murine macrophages across niches

**DOI:** 10.1101/2024.12.20.629843

**Authors:** Cassandra J. McGill, Olivia S. White, Ryan J. Lu, Nirmal K. Sampathkumar, Bérénice A. Benayoun

## Abstract

Macrophages are a key cell type of the innate immune system and are involved at all steps of inflammation: (i) they present antigens to initiate inflammation, (ii) they clear up foreign bodies through phagocytosis, and (iii) they resolve inflammation by removing or deactivating mediator cells. Many subtypes of macrophages have been identified, classified by their niche and/or embryonic origin. In order to better develop therapies for conditions with macrophage dysfunction, it is crucial to decipher potential sex-differences in key physiological mediators of inflammation so that treatment efficacy can be ensured regardless of biological sex. Here, we conduct a meta-analysis approach of transcriptomics datasets for male *vs*. female mouse macrophages across 8 niches to characterize conserved sex-dimorphic pathways in macrophages across origins and niches. For this purpose, we leveraged new and publicly available RNA-sequencing datasets from murine macrophages, preprocessed these datasets and filtered them based on objective QC criteria, and performed differential gene expression analysis using sex as the covariate of interest. Differentially expressed (DE) genes were compared across datasets and macrophage subsets, and functional enrichment analysis was performed to identify sex-specific functional differences. Consistent with their presence on the sex chromosomes, three genes were found to be differentially expressed across all datasets (i.e. *Xist*, *Eif2s3y*, and *Ddx3y*). More broadly, we found that female-biased pathways across macrophage niches are more consistent than male-biased pathways, specifically relating to components of the extracellular matrix. Our findings increase our understanding of transcriptional similarities across macrophage niches and underscore the importance of including sex as a biological variable in immune-related studies.

**Highlights:**

- Across 17 independently transcriptomic macrophage datasets, three genes are always differentially expressed between males and females: *Ddx3y*, *Eif2s3y*, and *Xist*.
- Jaccard analysis reveals female macrophages have more stereotypical transcriptional profile than male macrophages across niches.
- Functional enrichment analyses show that female-biased pathways across macrophage subtypes are enriched for extracellular matrix-related genes.

## Introduction

Macrophages are a key component of the mammalian immune system and are responsible for producing chemokines that activate the rest of the immune system to combat infection. They can be derived from circulating monocytes or from embryonic progenitors in the yolk sac and fetal liver, giving rise to tissue-resident macrophages^1^. After birth, the primary source of new macrophages comes from the differentiation of monocytes, which can be recruited to tissues throughout life. Macrophages are generally categorized by the tissues they are found in, which is supported by the importance of niche signals and of original progenitors on mature macrophage phenotype ^2–4^. Proper regulation of macrophage transcriptional landscapes is crucial for health and homeostasis ^5^.

The mammalian immune system shows significant differences between males and females in both innate and adaptive immune responses ^6–10^. Generally, females exhibit a more robust immune response, which may contribute to a higher risk of autoimmunity, whereas males have increased susceptibility to infections and poorer outcomes from sepsis ^11,12^. Despite the prevalence of sex differences in multiple immune-related diseases, the precise mechanisms underlying these differences are not yet fully understood. Macrophages exhibit sex-specific differences in their transcriptional profiles and functional states, which have significant implications for health and disease. Excessive macrophage activation is associated with numerous conditions, including neurodegeneration, atherosclerosis, osteoporosis, and cancer, many of which exhibit sex-biased tendencies. Understanding the sexually dimorphic nature of macrophages is crucial for advancing precision medicine and developing targeted therapies for immune-related diseases.

Despite the prevalence of sex differences in multiple immune-related diseases, the precise mechanisms underlying these differences are not yet fully understood. The “Immunological Genome Project” (ImmGen) consortium explored transcriptional profiles across male and female immune cells. Interestingly, they reported minimal sex-based differences in most immune cells, with the exception of macrophages from three distinct tissues—peritoneal cavity macrophages, spleen macrophages, and microglia. These macrophage subtypes exhibited pronounced sex-biased transcriptional patterns, suggesting that females may have a more immune-activated state ^6^. However, a comprehensive and systematic study of sex-differences across macrophage subtypes has not yet been performed.

Here, we leveraged new and publicly available transcriptome datasets from murine macrophages across a variety of niches. Although we initially identified 21 datasets for our study, only 18 datasets were retained after various quality filtering steps for downstream analysis. Intriguingly, we found little overlap of differentially expressed genes by sex across macrophage subtypes, with the only universally-shared DEGs being X-or Y-linked. We performed both over-representation analysis (ORA) and gene set enrichment analysis (GSEA) to identify sex-dimorphic pathways across macrophages subtypes. Contrary to the gene-level analysis, pathway-level analysis showed conserved sex-biased enrichments, with cell cycle-related pathways in male macrophages and enrichment of extracellular matrix (ECM) and immunoregulatory interaction pathways in female macrophages across multiple niches. Our analyses suggest that there are niche-specific effects of sex on the transcriptome of macrophages, and that the ECM may be particularly important to females’ immune response.

## Results

### Differential gene expression analysis finds multiple shared sex-dimorphic genes across various macrophage niches

We identified 21 datasets, new and publicly available (**Supplementary Table S1**), with sequencing-based transcriptomic profiling of male vs. female murine macrophages (**Figure 1A**). We started with 6 microglia, 5 peritoneal, 3 alveolar, 2 bone-marrow-derived macrophages (BMDM), 2 osteoclast precursor (OCP), 1 exudate, 1 spleen, and 1 pleural dataset. Macrophages can originate embryonically as yolk sac progenitors (microglia, lung macrophages), or postnatally from bone marrow-derived monocytes (BMDM, OCP). Interestingly, peritoneal macrophages originate from yolk sac progenitors, but are replenished during inflammation and aging with progenitors from the bone marrow^13^.

**Figure 1.**
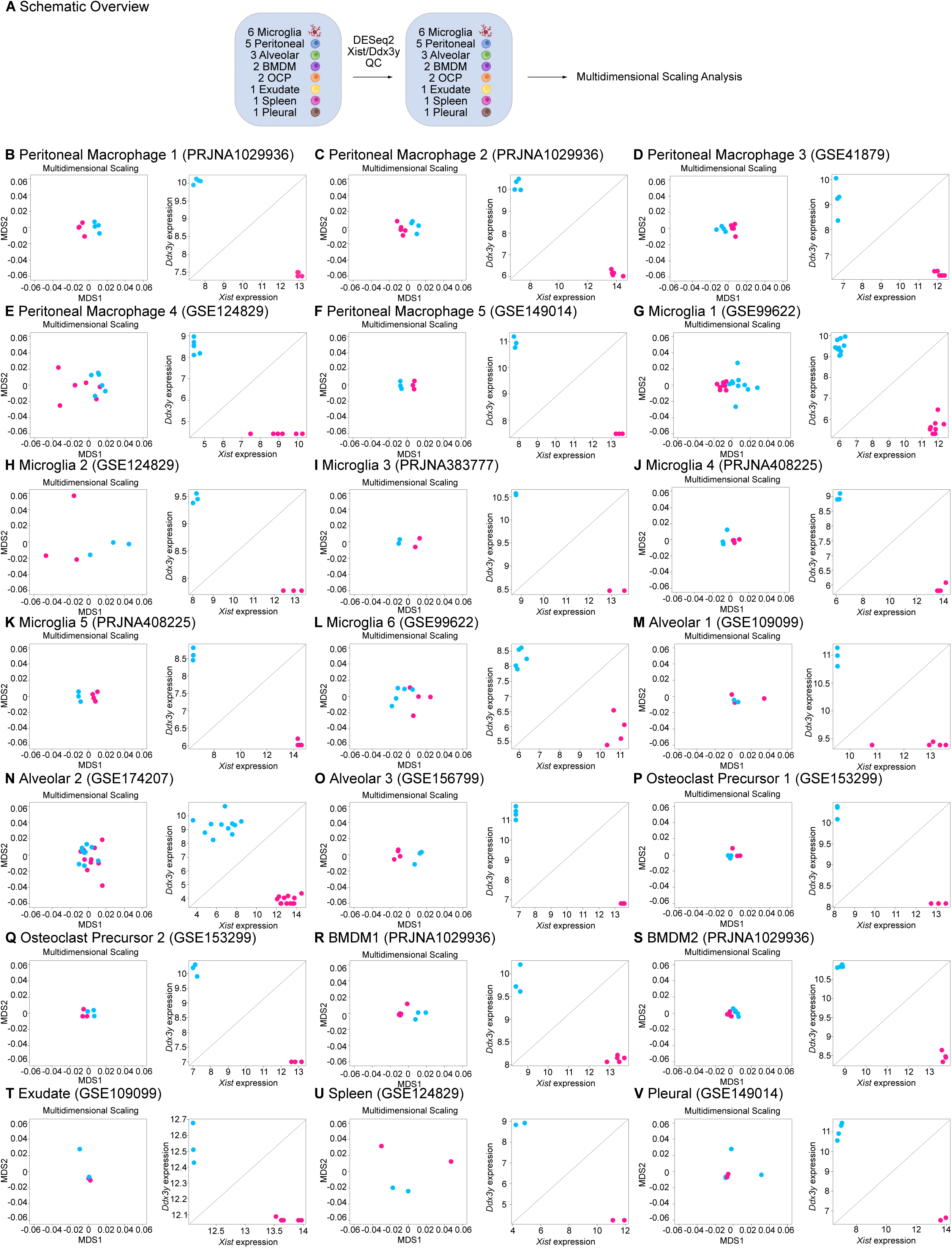
Quality Control of Bulk RNA-seq of Macrophage Niches. (A) Schematic overview of the experimental design. Multidimensional Scaling and Normalized log2(counts) *Xist* expression (X-axis) vs Normalized log2(counts) *Ddx3y* expression (Y-axis) for peritoneal macrophage datasets (B-F), microglia datasets (G-L), alveolar datasets (M-O), osteoclast precursor cells (P-Q), bone marrow derived macrophages (R-S), exudate dataset (T), spleen dataset (U), and pleural dataset (V). For all graphs, pink indicates female samples and blue indicates male samples.

After performing expression normalization using DESeq2 in R, expression levels of male-specific *Ddx3y* and female-specific *Xist* genes across all samples were examined to ensure proper labeling (**Figure 1B-V**). We also performed multi-dimensional scaling analysis (MDS) to determine whether sex could separate macrophage sample transcriptomes, which revealed clear separation across analyzed datasets (**Figure 1B-V**). Differentially expressed genes (DEGs) between males and females were compared within each dataset. One microglia, spleen, and pleural datasets with fewer than 50 DEGs were excluded. The remainder of this analysis was performed on five microglia, five peritoneal, three alveolar, two BMDMs, two OCPs, and one exudate dataset (**Figure 2A**). There was a wide range in the number of DEGs across datasets, with alveolar dataset 3 having 7130 DEGs and BMDM dataset 2 having only 51 (**Figure 2B**). This discrepancy may come from several factors, including (i) number of replicates in each study, (ii) absolute depth of sequencing for each study, (iii) spurious technical noise specific to each study.

**Figure 2.**
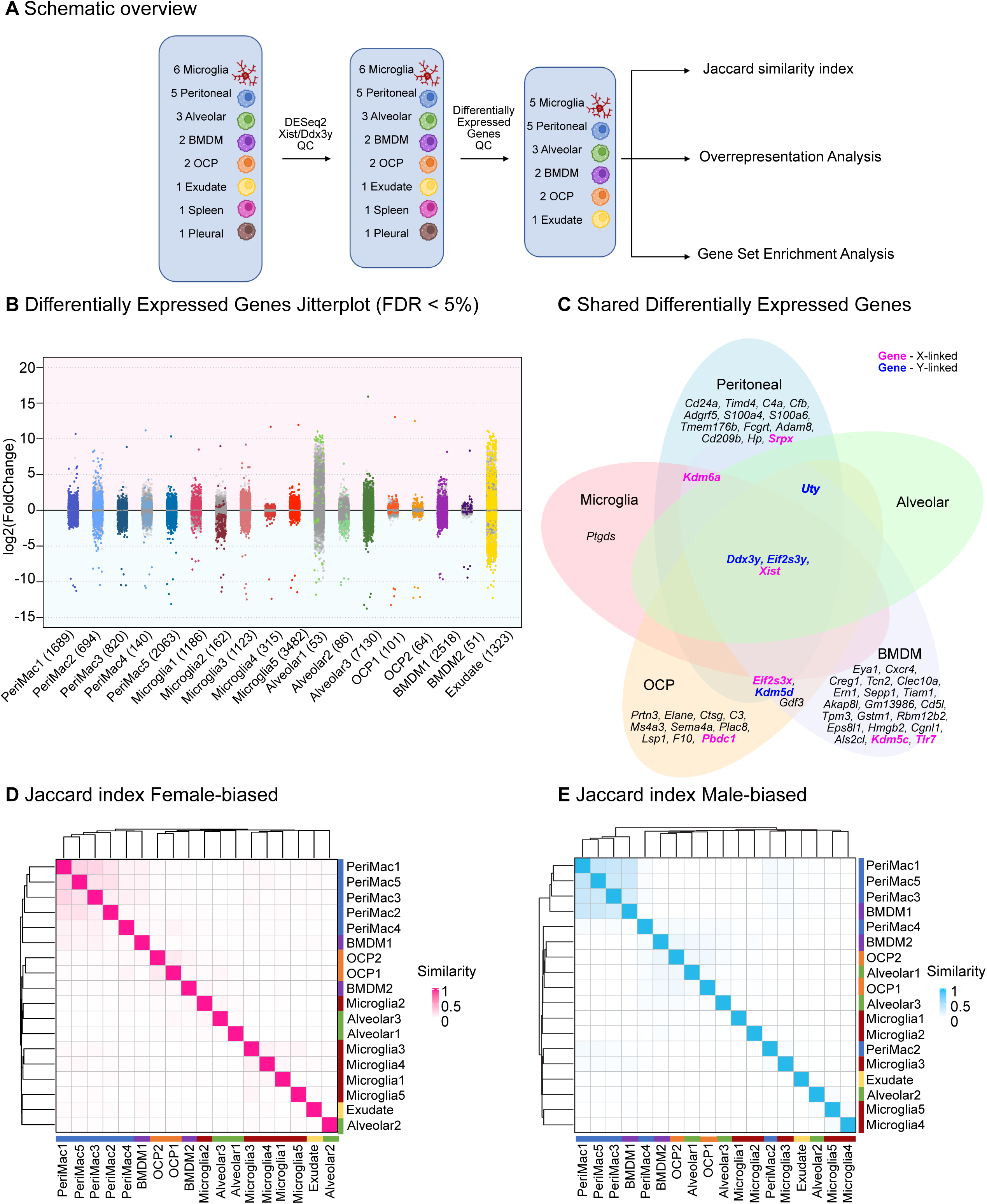
Differential Gene Expression across Macrophage Niches. (A) Schematic overview of the experimental design. (B) Jitterplot of differentially expressed genes between males and females across all datasets. Grey dots represent the log2 fold change values of all genes in the dataset, and dots with color are those that are significantly differentially expressed (FDR < 5%). Log2 fold change > 0 indicates female biased expression. Blue represents peritoneal macrophages (PeriMac), red represents microglia, green represents alveolar macrophages, orange represents osteoclast precursor cells (OCP), purple represents bone marrow-derived macrophages (BMDM), and yellow represents exudate macrophages. (C) Venn diagram of differentially expressed genes between macrophage niches. Genes with FDR of <5% were considered significantly differentially expressed. Datasets were combined into niches and the differentially expressed genes that each niche had in common were compared. Bold indicates sex chromosome-linked genes. Pink indicates X-linked and blue indicates Y-linked genes. (D) Female-biased jaccard index heatmap. Light coloring of the plot indicates overall low similarity between female-enriched genes across datasets. (E) Male-biased jaccard index heatmap. Light coloring of the plot indicates overall low similarity between male-enriched genes across datasets.

We then combined DE gene sets according to niches to compare identified DEGs. The peritoneal macrophages had 20 DEGs shared across all five datasets, the microglia had five shared DEGs across the five datasets, alveolar macrophages had 4 shared DEGs shared across 3 datasets, BMDMs had 28 shared DEGs across 2 datasets, OCPs had 17 shared DEGs across 2 datasets, and the singular exudate macrophage dataset had 1323 DEGs (**Figure 2B,C**). Three genes were found to be differentially expressed across all datasets: *Xist*, *Ddx3y*, and *Eif2s3y* (**Figure 2C**). *Xist*, also known as X-inactive specific transcript, is located on the X chromosome and primarily involved in X-chromosome dosage compensation, leading to the silencing of one of the two X chromosomes in females ^14^. *Ddx3y* (DEAD-box helicase 3) is a spermatogenic factor, and *Eif2s3y* (eukaryotic translation initiation factor 2, subunit 3) similarly regulates the proliferation of spermatogenial stem cells ^15^. Both genes are located on the Y chromosome. Genes shared amongst 2 or more datasets also tended to be sex-chromosome specific, including X-linked *Kdm6a* and *Eif2s3x*, and Y-linked *Uty* and *Kdm5d*. Interestingly, a DE gene identified as sex dimorphic in both OCPs and BMDMs, *Gdf3*, was not sex-chromosome linked (**Figure 2C**). *Gdf3*, also known as growth differentiation factor-3, encodes a protein involved in TGFB signaling and the regulation of bone and cartilage growth ^16^.

We next examined the genes differentially expressed by sex that were specific to each niche. Peritoneal macrophages had 13 niche-specific DE genes, including members of the complement system (*C4a, Cfb*), S100 family (*S100a4, S100a6*), and X-linked gene *Srpx* (**Figure 2C**). Microglia had one niche-specific sex-DE gene, *Ptgds*, which is an enzyme that catalyzes Prostaglandin D2 (PGD2) production. PGD2 enhances microglial inflammation and is higher in females 4/5 of the analyzed microglia datasets (**Figure 2C**; **Supplemental Table 2N-S**) ^17^. Three OCP-specific DE genes by sex include *Prtn3*, *Elane*, and *Ctsg*, all of which encode neutrophil serine proteases. These proteases are crucial in immune responses and play a role in histone H3 proteolytic cleavage that leads to monocyte-to-macrophage differentiation ^18^. In the bone marrow, granulocyte-macrophage progenitors can differentiate into “neutrophil-like” monocytes that express higher level of these proteases, consistent with the transcriptional profile we see in the OCPs resulting from sex-differences in the major progenitor for OCPs ^19^. For BMDMs, niche-specific sex DE genes include X-linked *Kdm5c* and *Tlr7*, as well as RNA-processing related genes *Eya1*, *Creg1* and *Rbm12b2* and DNA regulation-related genes *Akap8l*, *Hmgb2*. Alveolar datasets showed no sex DE genes specific to that niche.

Next, we asked if, regardless of significance threshold (which can be impacted by the differential statistical power of various datasets due to replicate numbers and technical noise related to library construction), there were global similarities in genes tending to be expressed in a female-or male-biased manner in macrophages across niches. For this purpose, we used the Jaccard index as a measure of similarity for the top 1500 sex-biased genes in all datasets and performed clustering analysis. Our results indicate that female-biased genes were more consistent among the cell types compared to male-biased genes (**Figures 2D,E**), with all the peritoneal macrophages clustering together, followed by bone-related macrophages, then microglia and lung macrophages.

### Overrepresentation analysis reveals shared sexually dimorphic pathways across macrophage niches

Next, we leveraged DE genes for each niche to perform over-representation analysis (ORA) for Gene Ontology terms. To improve our analytical power (and reflect biology), we used a list of DEGs shared across a majority of datasets in a niche (4/5 in peritoneal macrophages, 4/5 in microglia, 2/3 in alveolar, 2/2 in BMDM, 2/2 in OCP) as genes of interest. All expressed genes from all datasets within that niche were used as the background (**Figure 3A, 4A**).

**Figure 3.**
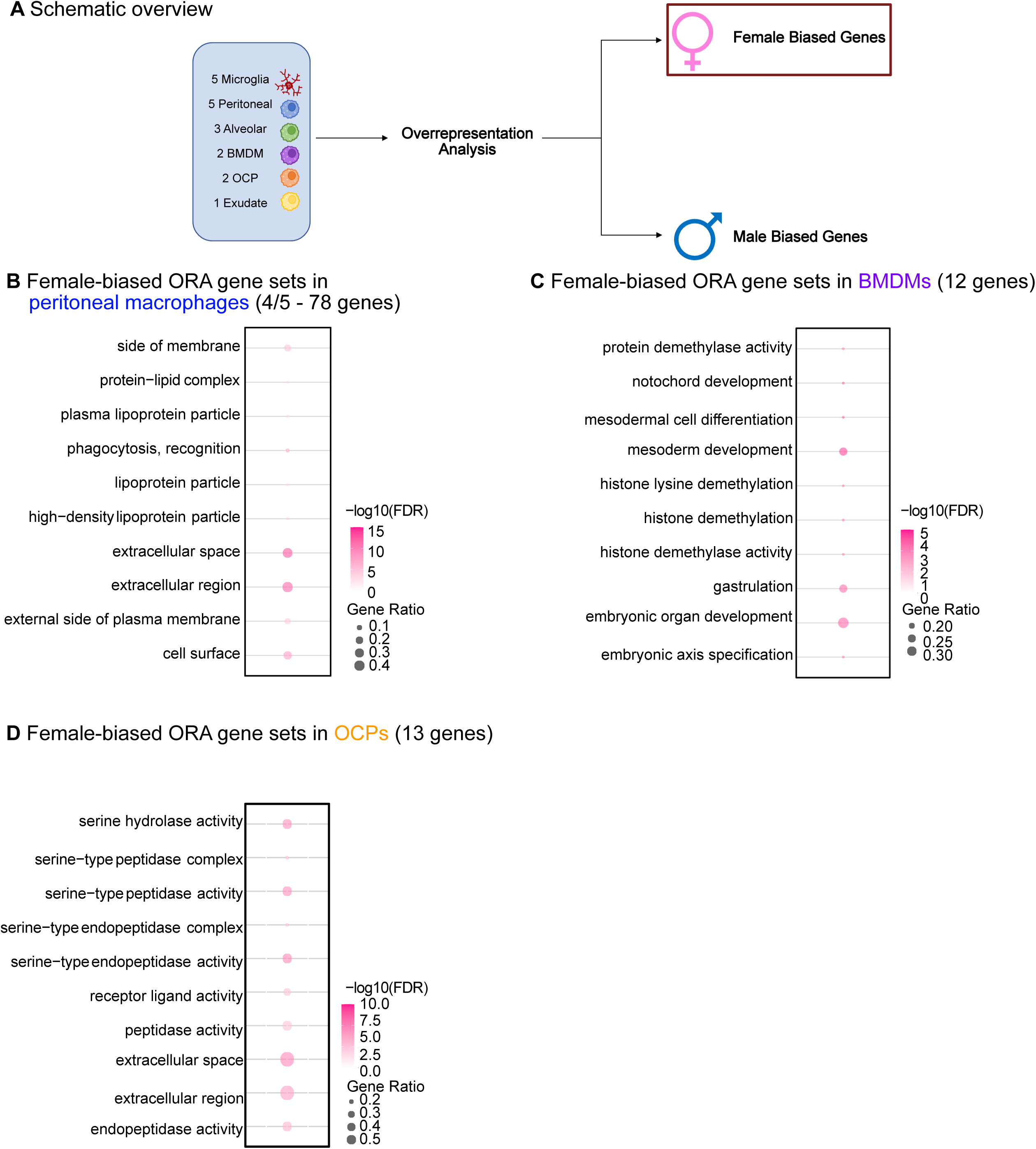
Gene Ontology overrepresentation analysis of macrophage female biased gene expression. (A) Schematic overview. The differentially expressed genes shared across all datasets in a niche were split into up-and down-regulated corresponding to female and male-biased genes. These were taken as the genes of interest and all expressed genes from all datasets within that niche were taken as the background. (B) Bubble plot of female-biased gene sets in 4/5 peritoneal macrophage datasets. (C) Bubble plot of female-biased gene-sets in BMDM datasets. (D) Bubble plot of female-biased gene-sets in OCPs.

In peritoneal macrophages, there were only significant GO terms (FDR < 5%) for female-biased DEGs (**Figure 3B, Supplementary Table S3B**). Among these include ‘external side of plasma membrane’, ‘extracellular space’, and ‘phagocytosis’ (**Figure 3B**). BMDMs showed both significant male and female biased terms, with females enriched for GO terms related to mesoderm development and embryonic organ development (**Figure 3D, Supplementary Table S3C**) and males enriched for nucleic acid binding and demethylation terms (**Figure 4C**, **Supplementary Table S3D**). OCPs showed both significant male and female biased terms, with female-biased genes showing enrichment for extracellular space (**Figure 3D**, **Supplementary Table S3E**), and males enriched for demethylation-related terms (**Figure 4C, Supplementary Table S3F**). In microglia, although no significant terms were observed for female-biased DEGs, male-biased DEGs were associated with GO terms related to nucleic acid binding, oxidoreductase activity, and dioxygenase activity (**Figure 4B, Supplementary Table S3A**). Exudate macrophages only had male-biased terms including sterol transfer activity and MHC protein complex binding (**Figure 4C, Supplementary Table S3G**). Alveolar macrophages also had only male biased GO terms, and these include protein demethylase activity and histone demethylase activity (**Supplementary Table S3H**).

**Figure 4.**
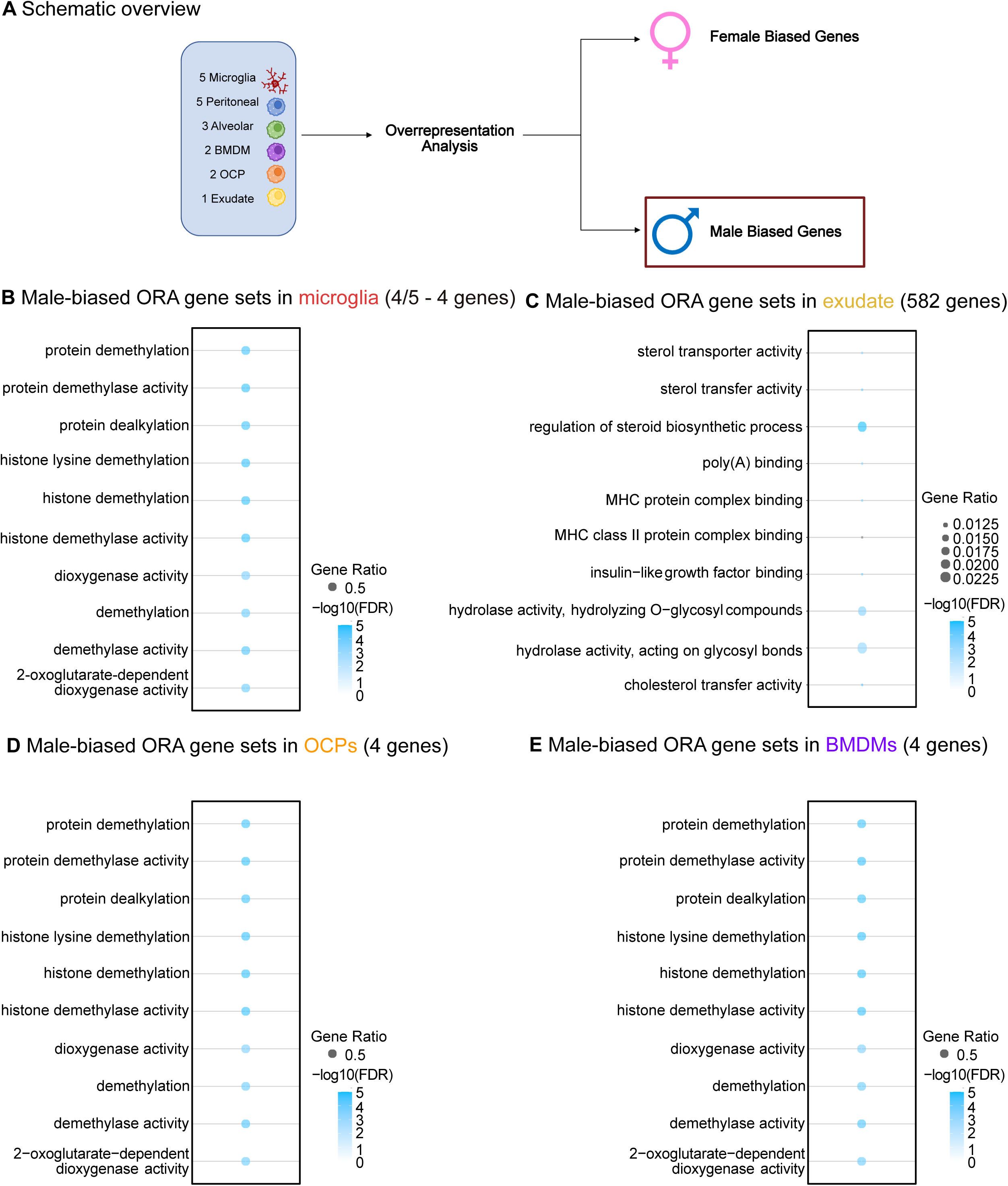
Gene Ontology overrepresentation analysis of macrophage male biased gene expression. (A) Schematic overview. The differentially expressed genes shared across all datasets in a niche were split into up-and down-regulated corresponding to female and male-biased genes. These were taken as the genes of interest and all expressed genes from all datasets within that niche were taken as the background. (B) Bubble plot of male-biased gene sets in 4/5 microglia datasets. (C) Bubble plot of male-biased gene-sets in the exudate dataset. (D) Bubble plot of male-biased gene-sets in OCP datasets. (E) Bubble-plot of male-biased gene-sets in BMDM datasets.

Interestingly, the ‘extracellular space’ GO term was found to be female-biased across both peritoneal macrophages and osteoclast precursor cells. To investigate this observation further, we thus performed GSEA using curated lists of Extra-Cellular Matrix (ECM)-related genes ^20^ (**Supplemental Figure S1A**). Consistent with our hypothesis, we found that, across niches, there is a significant female bias of ECM-related gene set expression in most peritoneal macrophages and OCP datasets (**Supplemental Figure S1A, Supplementary Table S5**). Interestingly, these observations are reminiscent of our previous work where we also observed a female-bias in the expression of ECM-related genes in another key mouse innate immune cell type, neutrophils ^21,22^. This suggests that female-bias in ECM-related gene expression may be a more general phenomenon across multiple immune cell types.

The ‘nucleic acid binding’ GO term was male-biased across microglia, BMDMs, and OCPs (**Figure 4B, D, E**). To note, genes with male-biased expression amongst all three macrophage types and significant GO terms include *Uty* and *Kdm5d* (both located on the Y chromosome). While shared terms in males across niches relate primarily to the sex chromosome, shared terms in females relate more to function and ECM biology.

### Gene set enrichment analysis finds similarly divergent pathways across all macrophage niches

We then leveraged these individual gene sets to perform Gene Set Enrichment Analysis (GSEA), which reduces the impact of threshold effects and highlights the impact of consistent changes across gene sets, using both Gene Ontology and REACTOME databases (**Figure 5A**). We found strong similarities amongst the datasets, with male-biased genes showing overall enrichment for GO terms relating to demethylation and T cell antigen processing and female-biased genes showing overall enrichment for GO terms relating to cell surface and dosage compensation (**Figure 5B**). In addition, GSEA analysis of REACTOME gene sets showed male-biased expression for cell cycle related pathways, and female-biased expression for GPCR signaling and extracellular matrix organization (**Figure 5C**; consistent with our ECM observation, **Supplemental Figure S1A**).

**Figure 5.**
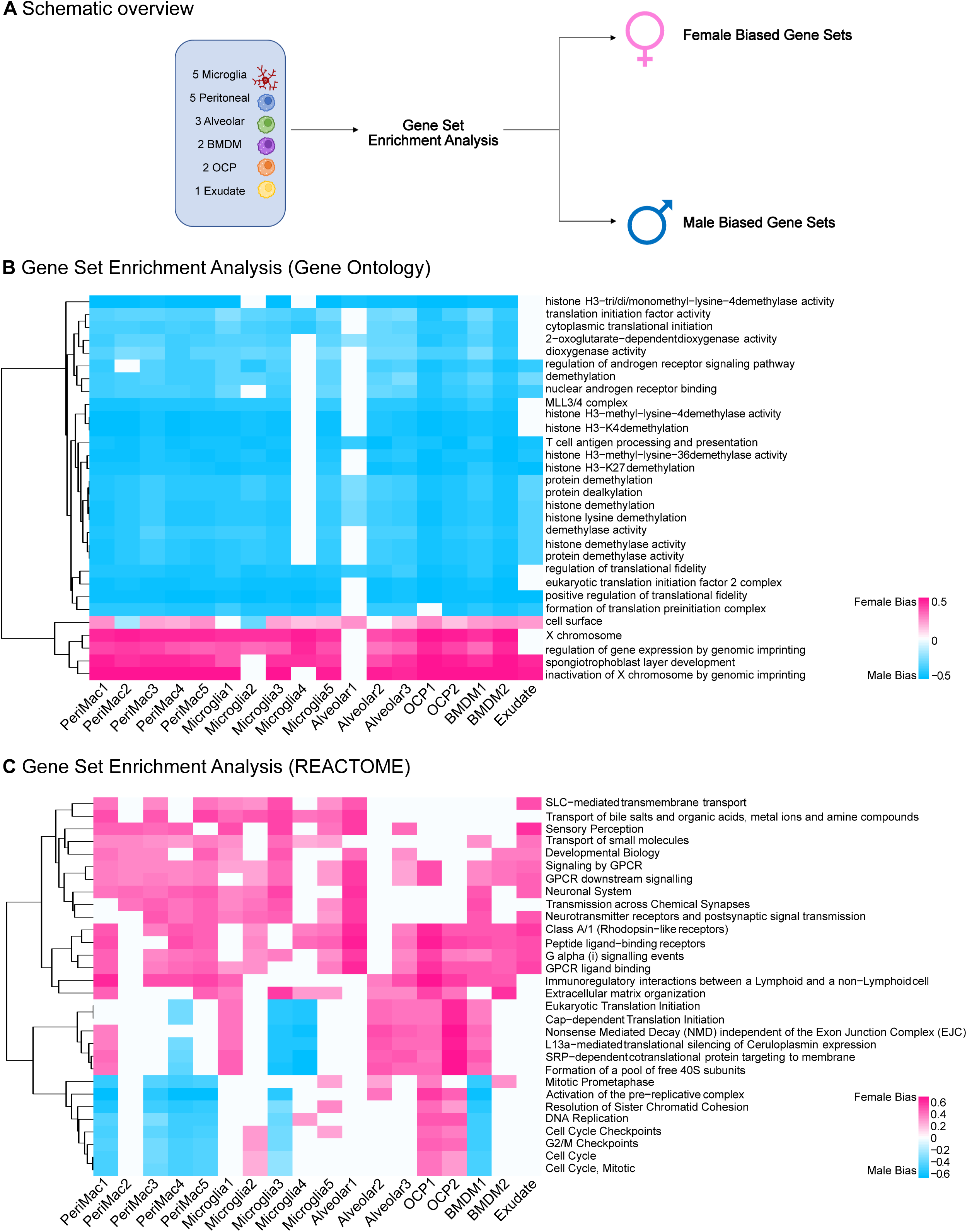
Analysis of recurrently sex-biased gene sets in murine macrophages from Gene Ontology and Reactome by Gene Set Enrichment Analysis. (A) Schematic overview. Gene set enrichment analysis was performed on each dataset using both Gene Ontology (GO) and REACTOME gene sets. (B) Heatmap of top 30 gene ontology terms significant in each dataset (FDR5%). (C) Heatmap of top 30 REACTOME pathways significant in each dataset (FDR5%). Color intensity indicates the level of enrichment: pink denotes enrichment in females and blue indicates enrichment in males.

Together, our analysis of the shared gene sets and pathways across all datasets suggests sex differences in ECM organization among macrophage subtypes.

## Discussion

Tissue-resident macrophages are crucial to proper immune system function. While all macrophages share the responsibility of clearing cellular debris and foreign bodies, tissue-resident macrophages also have unique responsibilities that facilitate homeostasis throughout the body. Microglia are responsible for preserving damaged myelin sheaths, osteoclasts facilitate resorption of bone, and alveolar macrophages recycle excess surfactant from the lungs ^23^. Sex differences have previously been reported in macrophages, with female macrophages having higher phagocytic activity than males ^24^.

Indeed, sex differences in immune responses have been well characterized with females exhibiting higher rates of autoimmune diseases. Females are known to mount higher antibody responses across many vaccines ^25^. However, the questions of why and how these sex differences occur have yet to be fully answered.

While individual studies have looked at sex differences within one macrophage niche ^26–29^, little work has been done to compare across macrophage niches ^6^. Here, we investigated the hypothesis that there is shared sex-dimorphic gene expression across macrophage niches. Our findings demonstrate that male macrophages, despite niche/origin, only consistently show differences in expression of Y-linked genes compared to niche-matched female macrophages. In contrast, we find that female macrophages, most specifically peritoneal and OCPs, in addition to X-linked genes, show increased expression of ECM genes compared to niche-matched male macrophages. Collectively, our findings suggest that there are consistent sex differences amongst multiple macrophage niches, and that female peritoneal macrophages and OCPs may remodel their ECM differently than male macrophages, which may contribute to phenotypes such as enhanced wound healing or even fibrosis^30–32^.

Importantly, the development of efficacious therapeutic interventions for immune/inflammatory disorders relies heavily on understanding macrophage biology in addition to sex-specific responses. For example, Alzheimer’s disease (AD) is heavily associated with microglia and mutations in macrophage-expressed genes ^33^. AD is also known to both present and progress differently by sex, indicating underlying mechanisms in microglia that may be sex dimorphic and create these different disease presentations ^34^. A group of cancer-like disorders called histiocytosis are the only known diseases to arise from somatic activating mutations in macrophages ^33^, and epidemiological studies indicate that these diseases are more prevalent in males than females ^35^. Dysfunctional OCPs are associated with development of osteoporosis, a disease that is four times more prevalent in women ^36^. Thus, investigating female and male-biased processes in macrophages, including the contribution of the ECM, will be an important step in developing treatments for diseases including, but not limited to, AD, histiocytosis, and osteoporosis.

## Methods

### Mouse husbandry

All animals were housed and treated in accordance with the Guide for Care and Use of Laboratory Animals. All experimental procedures were approved by the Institutional Animal Care and Use Committee (IACUC) at the University of Southern California, in accordance with institutional and national guidelines. Samples were derived from animals approved on IACUC protocol #20770.

### Data acquisition

Publicly available datasets were identified using (i) literature search for manuscripts describing RNA-seq of macrophages with female and male samples, and (ii) search on GEO and SRA. These datasets are listed in **Supplementary Table S1**. All data was downloaded and processed using a unified preprocessing pipeline.

Four new datasets were generated for this study using macrophages from 4 months old C57BL6/JNia mice – two independent datasets from isolated naïve peritoneal macrophages (Peritoneal Cohort 1, and Peritoneal Cohort 2) and two independent datasets from bone-marrow derived macrophages (BMDM Cohort 1, and BMDM Cohort 2). The raw sequencing data was deposited to the Sequencing Read Archive (SRA) under accession number PRJNA1029936. Isolation of primary macrophages and ‘omic’ library construction are described below.

### Isolation of mouse peritoneal macrophages

To isolate peritoneal macrophages, 10mL of 3% PBS/BSA was carefully injected into the intraperitoneal cavity, and the injected PBS/BSA was recovered to obtain naive peritoneal cells. The peritoneal lavage was centrifuged at 300g for 10 minutes at 4°C. Samples were then treated with 1x red blood cell lysis buffer (Miltenyi Biotec, no.130-094-183) for 2 minutes at room temperature, followed by centrifugation at 300g for 10 minutes at 4°C, to remove any potential red blood cells. The samples were then passed through a 30µM MACS Smartstrainer (Miltenyi-Biotec, no.130-110-915) to obtain a single cell suspension. Peritoneal macrophages were then isolated using the mouse peritoneal macrophage solation Kit (Miltenyi Biotec, no.130-110-434) according to the manufacturer’s instructions. Isolated cells were pelleted and snap frozen prior to RNA extraction (see below).

### Isolation of macrophages from mouse bone marrow

Long bones (tibia and femur) of young and aged mice from both sexes were collected and bone marrow was flushed into 1.5mL Eppendorf tubes via centrifugation (30 seconds, 10,000g) ^37^. The recovered bone marrow was treated with 1x red blood cell lysis buffer (Miltenyi Biotec, no.130-094-183) for 2 minutes, followed by centrifugation at 300g for 10 minutes at 4°C. After resuspension, cells were filtered on a 70µM filter (Miltenyi Biotec, no. 130-110-916), and pelleted again to remove any debris. Cells were then cultured in DMEM/F12 (Corning, no. 10-090), 10% FBS (Millipore Sigma, no. F0926), 10% L929 (ATCC CCL-1) conditioned media (Collected after 48 hours in T175 flask (VWR, no. 10861- 576) and filtered with 0.45µM PVDF (VWR, no. 76010-408) ^38^), 1% Penicillin Streptomycin (Corning, no. 30-002-CI), and 2ng/mL recombinant M-CSF (Miltenyi Biotec, no. 130-101-706). To remove dead and floating cells, a media change was performed 72 hours after initial plating. After seven days (to allow for full differentiation into macrophages), cells were collected, pelleted and snap frozen prior to RNA extraction (see below).

### RNA extraction and RNA-seq library preparation

Frozen cell pellets were resuspended in 500µL of Trizol reagent (Thermo-Fisher, no. 15596026). Total RNA was purified using the Direct-zol RNA Kit (Zymo, no. R2052) following the manufacturer’s instructions. RNA quality and integrity was assessed using the 4200 TapeStation system (Agilent Technologies) with a High Sensitivity RNA ScreenTape (Agilent, no. 5067-5579). We used 450ng of total RNA to build RNA-seq libraries.

Total RNA was subjected to ribosomal-RNA depletion via the NEBNext rRNA Depletion Kit (New England Biolabs, no. E6310) followed by library preparation via the SMARTer Stranded RNA-seq Kit (Clontech-Takara, no. 634411), both according to the manufacturer’s instructions. Library quality was assessed using the 4200 TapeStation system (Agilent Technologies) with a High Sensitivity DNA ScreenTape 1000 (Agilent, no. 5067-5584).

Libraries were multiplexed for sequencing as 150bp paired-end reads on the Illumina Novaseq 6000 platform at Novogene (USA).

### RNA-seq analysis pipeline

Paired-end 150-bp reads were hard-trimmed to 75 bp using Fastx Trimmer. Trimmed reads were mapped to the mm10 genome reference using STAR v.2.7.0e. Read counts were assigned to genes from the UCSC mm10 reference using subread v.2.0.2 ^39^.

Read counts were imported into R (version 4.2.2) to perform downstream analyses. Only genes with mapped reads in at least half of the RNA-seq libraries in individual datasets were considered to be expressed and retained for downstream analysis. R package ‘sva’ v.3.46.0 ^40^ was used to estimate surrogate variables and the removeBatchEffect function from ‘limma’ v.3.54.2 ^41^ was used to regress out the effects of computed surrogate variables from raw read counts.

The ‘DESeq2’ R package (DESeq2 v.1.38.3) was used for data normalization and differential gene expression analysis ^42^. Genes with FDRL<L5% were considered statistically significant and are reported in **Supplementary Table S2**.

### Quality control

Read depth was calculated for each sample across all datasets, then averaged by sex to confirm no substantial discrepancies between male and female samples that could bias differential expression analysis. After normalization with DESeq2, normalized expression of *Ddx3y* and *Xist* were examined to ensure proper labelling of males and females (see **Figure 1**). Lastly, differentially expressed genes (DEG) were determined with FDR cutoff < 5% and datasets with fewer than 50 differentially expressed genes were excluded from downstream analysis, since these datasets are likely underpowered. This DEG cutoff resulted in the removal of 3 datasets (microglia dataset # 6, spleen, and pleural).

### Dimensionality reduction

To perform multi-dimensional scaling (MDS) analysis ^43^, we used a distance metric between samples based on the Spearman’s rank correlation value (1-Rho), which was then provided to the core R command ‘cmdscale’. MDS analyses were visualized in plots and visually inspected to ensure that the females and males clustered appropriately (Figure 1).

### Jaccard index

To compute the similarity between DE genes across datasets, the Jaccard index was used. The top 1500 upregulated and downregulated genes from each dataset were used to evaluate the similarity of effect size and direction of gene regulation across datasets without issues from list size imbalance.

The computed Jaccard index was then used to create clustered heatmaps of similarity across datasets.

### Functional enrichment analysis

Over Representation Analysis (ORA) was performed through the R package ‘clusterProfiler’ v4.6.2 ^44^, and Bioconductor annotation package ‘org.Mm.eg.db’ v3.16.0. The differentially expressed genes shared within each niche were divided into up and down-regulated based on the DEseq2 log fold change. These were used as the shared genes and all expressed genes across all datasets in that niche were used as the universe for the clusterProfiler function ‘enrichGO’. The ‘ggplot2’ v.3.5.1 package in R ^45^ was used to generate bubble plots for each niche. All significant results are in **Supplemental Table S3**.

The Gene Set Enrichment Analysis (GSEA) paradigm through its implementation in the R package ‘clusterProfiler’ v4.6.2 ^44^, and Bioconductor annotation package ‘org.Mm.eg.db’ v3.16.0 were used to perform the functional enrichment analysis. The DEseq2 t-statistic was used to generate the ranked list of genes for functional enrichment analysis.

Gene ontology (GO) terms (**Figure 4B**) or Reactome (‘ReactomePA’ v1.42.0) pathways (Figure 4C) were ranked by significance, and the top 30 for each are reported in Figure 4 ^46^. All significantly enriched gene sets are reported in **Supplementary Table S4.**

### Gene Set Enrichment Analysis of extracellular matrix-related terms

We obtained a manually curated list of gene sets relevant to extracellular matrix biology ^20^. We used the GSEA paradigm to determine whether ECM-related gene sets were differentially regulated as a function of sex in the datasets. For this purpose, we used R package ‘phenotest’ 1.38.0 in R version 4.0.2. The DEseq2 t-statistic for each gene was used to create the ranked gene list for functional enrichment analysis. The table output of this analysis is reported in **Supplementary Table S5** and graphically represented in **Supplemental Figure S1**.

## Acknowledgements

This work was supported by NIA T32 AG052374 and NIA F31 AG084279 predoctoral fellowship (C.J.M.); a generous gift from Kathleen Gilmore, Pew Biomedical Scholar award #00034120, and Hevolution HF-GRO-23-1199072-28 (B.A.B.).

## Author Contributions

B.A.B designed the study. R.J.L. and N.K.S. performed macrophage isolation and RNA-seq library construction. O.S.W. and C.J.M. performed data analyses. All authors contributed to the writing and editing of the manuscript.

## Data Availability

All newly generated data will be made available on GEO (Accession number PRJNA1029936). All other data is currently available in public databases, with accession numbers listed in **Supplemental Table 1A**.

## Code Availability

All analytical scripts for this study are available on the Benayoun lab github (https://github.com/BenayounLaboratory/ SMAC).

## Legend to Supplementary Figures

**Supplemental Figure S1.**
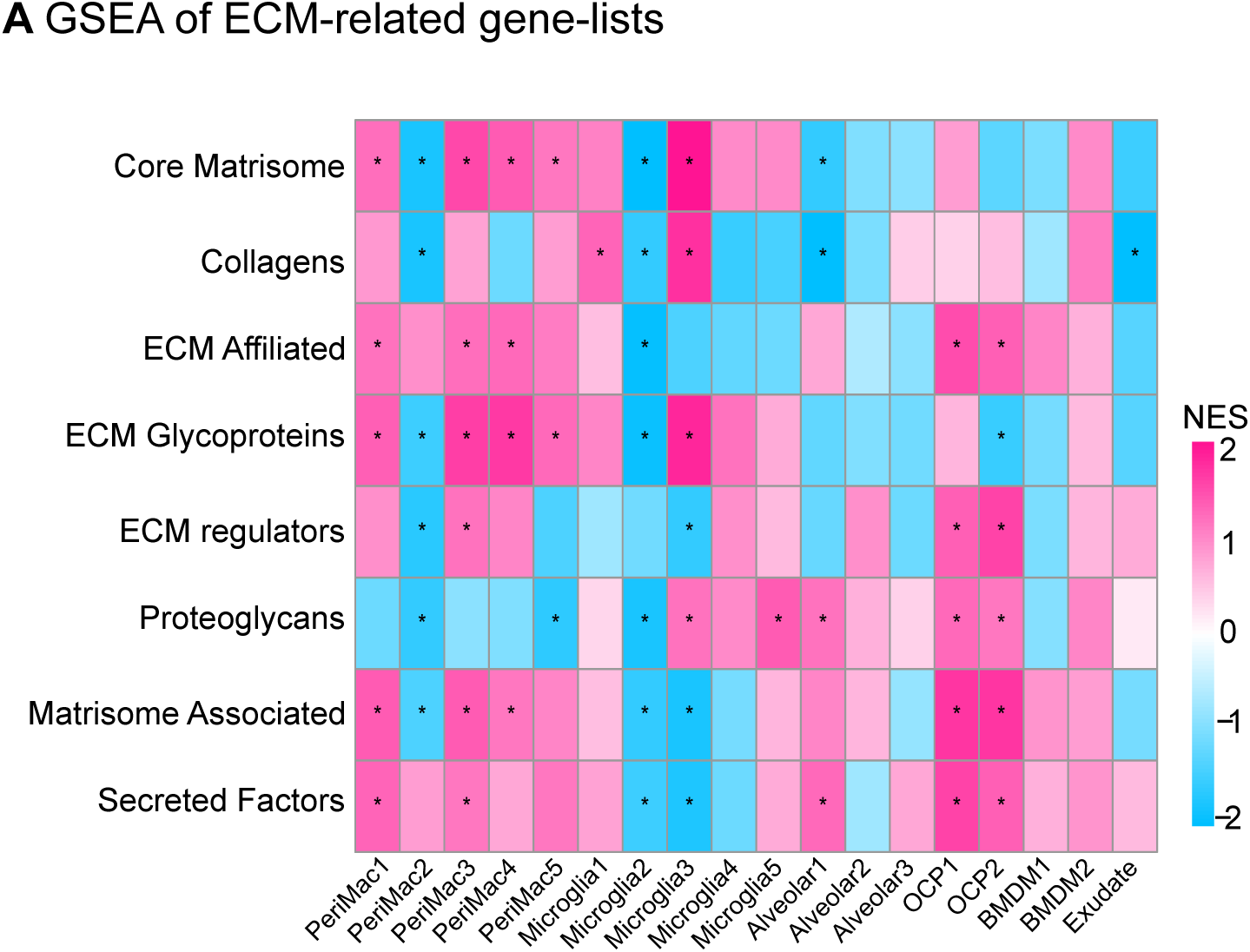
Gene Set Enrichment Analysis of Extracellular Matrix Gene Lists. (A) Heatmap showing GSEA enrichment results for ECM-related genes based on male-or female-biased expression from bulk RNA-seq data of various macrophage niches. Full analysis results are in **Supplemental Table 5**.

